# Multiple mating predicts queen reproductive output in the polygynous Neotropical trap-jaw ant, *Odontomachus chelifer* (Formicidae: Ponerinae)

**DOI:** 10.1101/2025.05.05.652256

**Authors:** Henrique Silva Florindo, Marianne Azevedo-Silva, Miguel Piovesana Pereira-Romeiro, Anete Pereira de Souza, Paulo S. Oliveira, Gustavo Maruyama Mori

**Affiliations:** Institute of Biosciences, São Paulo State University, São Vicente SP, Brazil; Department of Ecology and Evolutionary Biology, University of Michigan, Ann Arbor MI, USA; Department of Animal Biology, University of Campinas (UNICAMP), Campinas SP, Brazil; Department of Ecology, São Paulo University (USP), São Paulo, Brazil; Department of Plant Biology, University of Campinas (UNICAMP), Campinas SP, Brazil

**Keywords:** Reproductive sharing, polyandry, colony structure, fitness, Neotropics

## Abstract

The factors that influence the organisms’ ability to reproduce are diverse. In ants, social structure may contribute to queens’ Reproductive Output (RO), alongside environmental conditions and genetic attributes. Polygynous species like the Neotropical trap-jaw ant, *Odontomachus chelifer*, are valuable biological systems to investigate the contribution of these factors to the RO of queens. We hypothesize that *O. chelifer* queens’ RO increases with (1) higher multiple mating (polyandry) per queen; (2) higher queen heterozygosity; (3) greater food availability. We sampled workers from 18 colonies from the Brazilian Cerrado savanna (15 workers per colony) and used eight microsatellites to infer the number of male mates per queen, queens’ heterozygosity, and the proportion of offspring produced per queen. The latter was used to calculate the index of deviation from even offspring production (DEM) among queens. Arthropod communities from sampling sites were used as a proxy for food resources availability. We evaluated the relationships between DEM and the predictor variables using bivariate generalized linear models. Our results showed a positive relationship between polyandry and DEM (Coefficient = 0.41, P < 0.001). It suggests that the number of male mates likely benefits queen productivity, highlighting the role colony structure plays in the fitness of individual queens within *O. chelifer* colonies. Our findings did not support the remaining hypotheses. This work illuminates the role of social forms in evolutionary processes in ants, making way for similar studies in different social insects and contributes to a better understanding of the reproductive traits of trap-jaw ants.

## Introduction

Reproductive output (i.e. the number of offspring produced by an organism; hereafter RO), is a significant component of fitness (Orr, 2009). Thus, understanding the factors that drive differences in individual fitness is crucial to unveil ecological and genetic processes that influence species evolution. In eusocial insects like ants, queen’s RO can be influenced by a range of factors, such as number and identity of reproductive queens, colony’s genetic structure and interindividual relatedness, and environmental features (Hölldobler & Wilson, 1990; Steiner et al., 2009).

Polyandry, a social form in which a queen mates with multiple males, is known to increase intracolonial genetic diversity and may also offer direct benefits to the queen (Baer, 2016). Polyandrous queens are likely to increase their RO, longevity, and colony size, through increased sperm storage and lifespan (Baer, 2016; Crozier & Fjerdingstad, 2001; Fjerdingstad & Boomsma, 1998; Heinze & Schrempf, 2008; Ruttner et al., 1973). Parental genetic makeup, specifically heterozygosity and its relationship with fitness, has been a subject of extensive discussion (Chapman et al., 2009; Szulkin et al., 2010). In many cases, heterozygosity-fitness correlations (HFCs) are positive, but weak (Szulkin et al., 2010). In ants, while underrepresented in HFC studies, there is evidence that queen heterozygosity may be positively linked with higher colony-founding success (Vitikainen et al., 2015), queen lifespan and/or fecundity (Haag-Liautard et al., 2009). Thus, heterozygosity could improve queen RO, especially because ants tend to have small effective population sizes (Seppä, 2008). Also, environmental features, such as food availability and adequate nesting conditions play an important role in offspring production (Blüthgen & Feldhaar, 2009; Byk & Del-Claro, 2011). Depending on the timing of food supplementation, colonies with higher food input may increase RO (Deslippe & Savolainen, 1994), production of alates, and females:males ratio (Ode & Rissing, 2002). The study of how these factors may influence components of fitness remains a fertile ground for empirical work, notably considering regions with high levels of biodiversity, like the Neotropics.

## Study species

The Neotropical ant *Odontomachus chelifer* (Latreille, 1802) (Formicidae: Ponerinae) is a valuable biological system to investigate the factors that influence RO. The species nests on the ground and its diet is composed mostly of epigeic arthropods (Raimundo et al., 2009; Christianini & Oliveira, 2010; Bottcher & Oliveira, 2014). Colonies of *O. chelifer* are functionally polygynous and present hierarchical reproductive dominance among queens mediated by ritualistic antagonistic behavior, which result in highly-ranked queens laying more eggs and engaging less frequently in foraging activities outside the nest (Medeiros et al., 1992). Within-colony genetic diversity data revealed that *O. chelifer* queens may also be polyandrous with low average levels of mating frequency (Azevedo-Silva et al., 2023). However, the influence of food availability, polyandry and heterozygosity on queen RO remains to be addressed. Here, we intend to fill this gap by simultaneously investigating the role of these three factors on *O. chelifer* queen RO. We hypothesized that RO increases with the increase in (1) the level of polyandry; (2) the queen heterozygosity; (3) food availability.

## Materials and Methods

The data used in this work was obtained from Salles et al., (2018) and Azevedo-Silva et al. (2023). Three independent transects were established in a Cerrado reserve, located in the state of São Paulo, Southeast Brazil (22°18S, 47°11’W). Transects were 1300 ± 282 m (mean ± SD) distant from one another and were further subdivided into fragment edge (< 15 m from the fragment edge) and fragment interior (> 45 m from the fragment edge). In each transect, 15 workers from six colonies of *O. chelifer* were sampled after actively searching and identifying each colony’s entrance. *Odontomachus chelifer* diet is mostly carnivorous, including a broad range of arthropod taxa with well-defined prey preferences regardless of their abundance on the forest ground (Raimundo et al., 2009). From each transect, we used eight pitfall traps spaced by ∼40 m to sample epigeic arthropod communities. Ants were identified to the species or morphospecies level, while other arthropods were identified at the order level (Salles et al., 2018). We calculated Shannon diversity index for arthropod community using vegan 2.6-4 package (Oksanen et al., 2022) and estimated mean arthropod abundance for the edge and interior of each transect as operational variables for food availability.

Workers’ DNA were isolated and genotyped for eight species-specific polymorphic microsatellites (Azevedo-Silva et al., 2023; Lemos et al., 2020) using an automatic sequencer (ABI 3500, Applied Biosystems). Parental genotypes were reconstructed using COLONY 2.0 (Jones & Wang, 2010), which allowed the quantification of the number of queens per colony (*nq*), the number of male mates per queen and the number of offspring (workers) per queen (Azevedo-Silva et al., 2023). Based on the 81 inferred queens’ genotypes, we estimated their heterozygosity using the GENHET function (Coulon, 2010), which provides five estimates of individual heterozygosity. Given the correlation between the estimates (Figure S1), we used the proportion of heterozygous loci per individual (PHt) for downstream analyses. We divided the number of offspring per queen by the number of sampled workers per nest to obtain the proportion of workers produced per queen (*nfp*). We then used *nfp* to calculate an index of deviation from even offspring production among queens (DEM), which was used as a proxy for queen RO (Equation 1). The index indicates, proportionally, the deviation that the queen had from equal reproductive partition (i.e. a queen with an index of 1.3 has produced 30% more offspring than what would be expected if offspring production was even among queens in each colony). A value of 1 indicates that the queen’s offspring production was equal to the average for that colony.

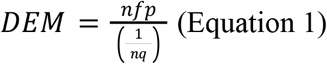

To test our hypotheses, we constructed four bivariate generalized linear models (GLMs) with gaussian distribution to evaluate queen RO based on our initial hypotheses using the *glm* function in R *stats* package (R Core Team, 2022), with the following predictor variables: (1) number of male mates; (2) proportion of heterozygous loci per queen; and food resource assessed by (3) arthropod diversity; (4) arthropod abundance; and (5) the interaction between arthropod diversity and abundance. Additionally, we considered that polyandry and queen heterozygosity effects on queen RO might overlap, as both variables contribute to offspring genetic diversity. To disentangle the potentially confounding effects of polyandry and queen heterozygosity on queens’ RO, we constructed a separate bivariate model restricted to monandrous queens only, using the proportion of heterozygous loci as the predictor variable. Otherwise specified, all analyses were conducted in R (R Core Team, 2022).

## Results and Discussion

The bivariate models indicated a significant positive association between polyandry and DEM (Coefficient = 0.41, SE = 0.0604, P < 0.001; Figure 1A), whereas the other variables did not present significant associations (Figure 1B-E). Our findings provide evidence that polyandrous *O. chelifer* queens have a higher reproductive performance compared to their nestmates, often presenting an above-average offspring production within the colony. Conversely, neither queen heterozygosity nor food availability were found to influence queen RO. One possibility is that chosen operational variables (i.e. arthropod abundance and diversity, and their interaction) may not have fully encompassed the concept of food availability for *O. chelifer*, due to the species’ well-defined prey preferences (Raimundo et al., 2009). Also, fallen fruits and seeds are additional food sources for *O. chelifer* that were not considered in this study, and that could contribute to colony fitness by providing specific nutrients (Bottcher & Oliveira 2014). While measuring the availability of such resources *in situ* presents a difficult task, controlled experimental studies could aid in informing the role of feeding habits on queen RO.

**Figure 1.**
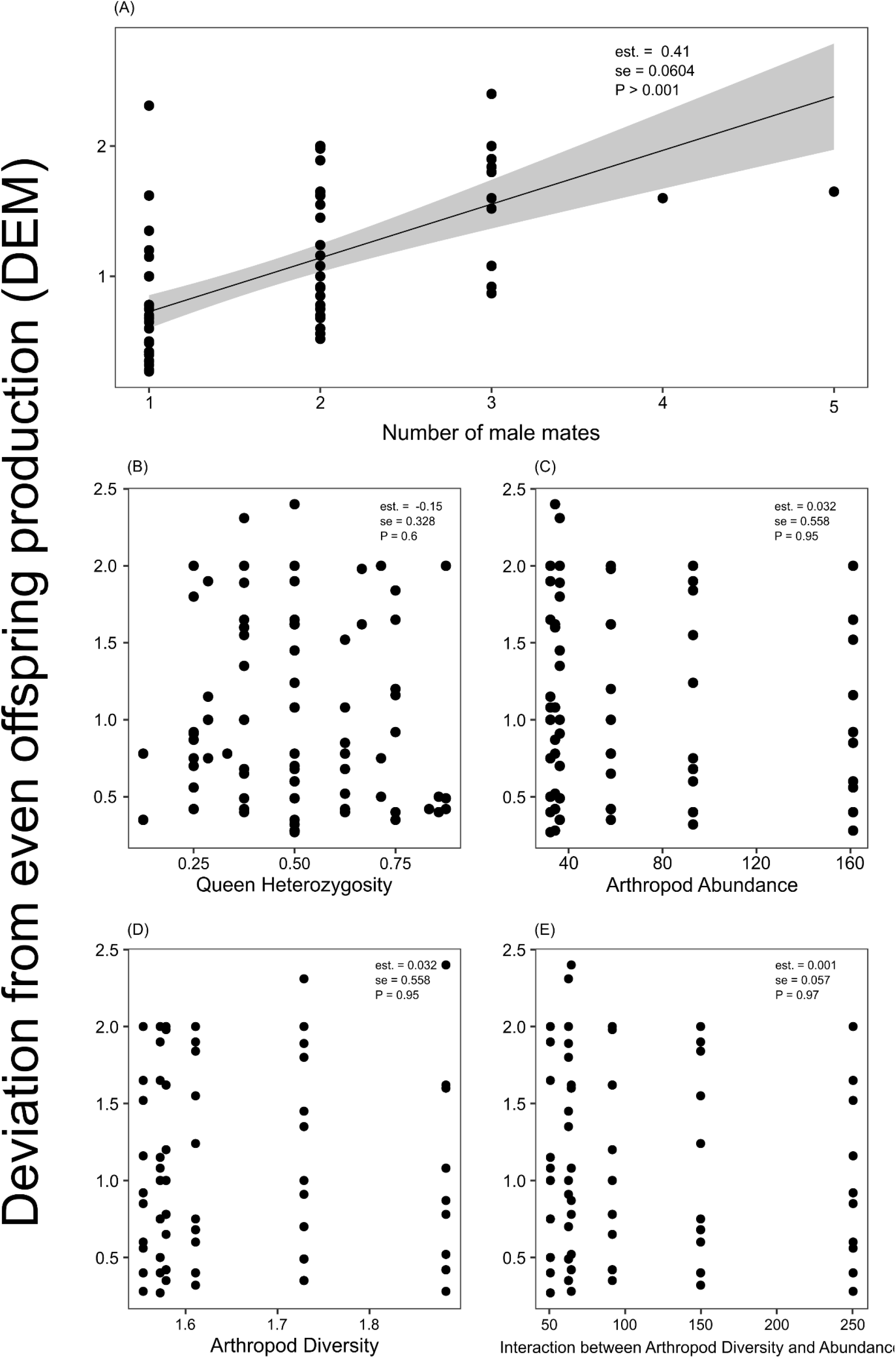
Bivariate analyses showing the relationships between different predictors and the index of deviation from even offspring production (DEM), used as a proxy of reproductive output of *Odontomachus chelifer* queens. Black circles indicate observed individuals, the line and highlighted area represent the model tendency and 95% confidence interval, respectively. **est** = coefficient estimate, **se** = standard error, **P** = P-value.

We recognize that the effects of individual heterozygosity could be concealed by the effects of polyandry on queen RO (given their similar contributions to intracolonial genetic diversity). However, we also found no significant association between heterozygosity and queen RO in the bivariate model accounting only for monandrous queens (Figure S3). While our findings offer no evidence of queen heterozygosity benefiting queen RO, we acknowledge that we may not be able to fully tease apart the effects of polyandry and heterozygosity on queen RO, as both variables were inferred from workers’ genotypes. Further studies investigating the impacts of heterozygosity among queens may allow us to better understand its effect on reproductive behavior, especially in scenarios where queen genetic makeup is the main driver of intracolonial genetic diversity, such as in colonies founded by individual queens.

Polyandry is found in at least one third of eusocial insect species (Boomsma & Ratnieks, 1996). Because the mating behavior of *O. chelifer* remains to be described, we argue that, despite initial costs such as additional effort to find mates and increased risk of infections (Baer, 2016; Hughes et al., 2008), polyandry may offer a range of benefits to the colony resulting from increased colony genetic diversity (Hughes et al., 2008). Such benefits may include pathogen resistance (Delaplane et al., 2021), more effective division of labor and brood rearing (Arnqvist & Nilsson, 2000; Baer, 2016), and increased capacity for colonies to adapt to environmental disturbance (Oldroyd & Fewell, 2007). For queens, polyandry is associated with an increased amount of sperm storage (Delaplane et al., 2015; Fjerdingstad & Boomsma, 1998), allowing them to produce more offspring over time (Boomsma & Ratnieks, 1996; Cole, 1983). Although the sperm limitation hypothesis is not a consensus for all eusocial species (Crozier & Page, 1985; Ruttner et al., 1973), theoretical work suggests a positive correlation between polyandry and colony size in monogynous colonies (Boomsma & Ratnieks, 1996). Since *O. chelifer* colonies are polygynous, it is possible that, due to the queens’ low mating frequency (Azevedo-Silva et al., 2023), a slight increase in sperm storage provided by few additional mates can increase queens’ offspring production.

Whether through fertility or queen physiology and lifespan, our findings indicate that polyandry provides benefits that translate into competitive advantages for queens (R. Crozier & Fjerdingstad, 2001; Fjerdingstad & Boomsma, 1998; Heinze & Schrempf, 2008). Because inseminated and more aggressive *O. chelifer* queens occupy the top of the colony’s dominance structure (Medeiros et al., 1992), our results suggest that highly-ranked queens (and consequently, with the greatest RO) could also have mated with more males. However, the possibility that multiple-mated queens of *O. chelifer* would also exert dominance over single-mated nestmates awaits further investigation.

In conclusion, we showed that polyandry plays a central role in the RO of *O. chelifer* queens, adding new information to the reproductive ecology and natural history in this species. Thus, our results suggest that not only the ritualistic antagonistic behavior among queens account for offspring partitioning, but also their reproductive behavior prior to colony foundation or acceptance. Our data adds to the understanding of how social traits are associated with ecological and evolutionary processes in ants.

## Conflict of interest statement

The authors have no conflict of interest.

## Data availability statement

The script and datasets used in the present study are available on author’s github (https://github.com/hsflorindo).

## Acknowledgements

We are grateful to Dr. Peter Nonacs, Luís F. P. Salles, and Alexander V. Christianini, for their insightful feedback and constructive suggestions during the final stages of this study. This work was supported by Coordenação de Aperfeiçoamento de Pessoal de Nível Superior (CAPES, Financial Code 001), Fundação de Amparo à Pesquisa do Estado de São Paulo (FAPESP 2017/18291-2; 2017/16645-1; 2020/15636-1; 2022/06529-2; 2022/02804-9) and Conselho Nacional de Desenvolvimento Científico e Tecnológico (CNPq 167161/2017-2; PIBIC 2020/1518; PIBIC 2021/4082).

## Authors contributions

The study was conceptualized by MAS, PSO and GMM. Genetic analyses were performed by MAS, HSF and MPR. HSF, MAS and MPR conceptualized the models and analyzed the data. HSF, MAS and GMM co-wrote the original draft. All authors took part in reviewing, providing feedback and granted approval for publication.

## Notes

### Competing Interest Statement

The authors have declared no competing interest.

