## Supplementary Material for "Multiple mating predicts queen reproductive output in the polygynous Neotropical trap-jaw ant, *Odontomachus chelifer* (Formicidae: Ponerinae)"

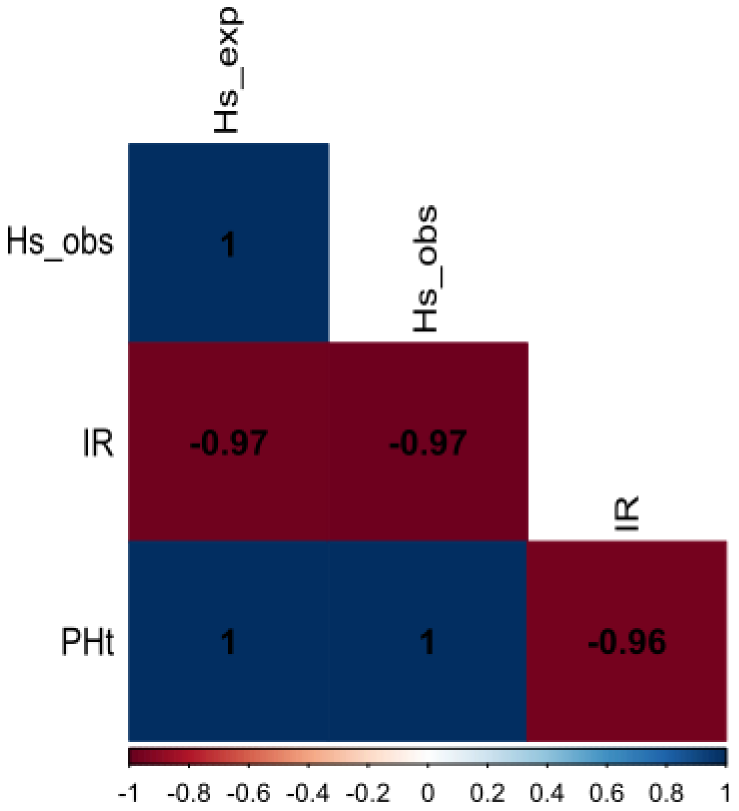


**Figure S1**: Correlation matrix for the heterozygosity estimates from the GENHET function; Hs_obs = Standardized heterozygosity based on the mean observed heterozygosity, IR = internal relatedness, PHt = proportion of heterozygous loci, HS_exp = Standardized heterozygosity based on the mean expected heterozygosity

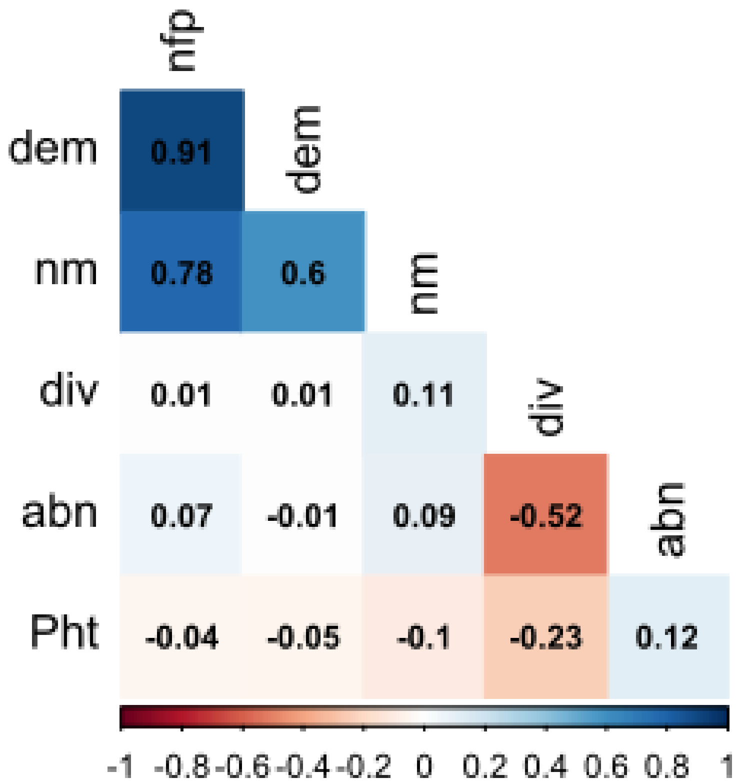


**Figure S2:** Correlation matrix including every variable used in the modeling process; nfp = proportion of offspring per queen, dem = deviation from even offspring production among queens, abn = food abundance, div = food diversity, Pht = proportion of heterozygous loci, nm = number of male mates per queen

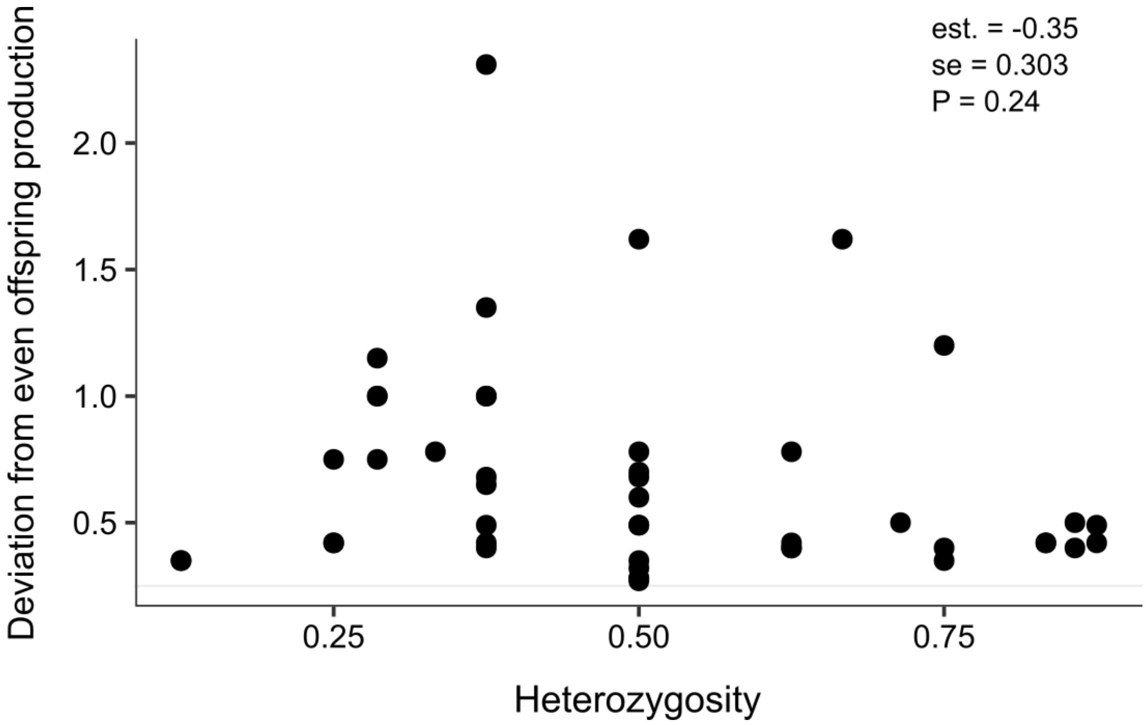


**Figure S3:** Generalized linear model showing the association between queen heterozygosity and the index of deviation from even offspring production for monandrous *Odontomachus chelifer* queens. Black circles indicate observed individuals. est. = coefficient estimate, se = standard error, P = P-value.
